# OPTIMOS PRIME: An R package for autoecological (optima and tolerance range) data calculation

**DOI:** 10.1101/654152

**Authors:** María Belén Sathicq, María Mercedes Nicolosi Gelis, Joaquín Cochero

## Abstract

1. Calculation of autoecological data, such as optima and tolerance ranges to environmental variables, can be useful to establish the distribution and abundance of the species. These calculations, although mathematically not complex, can be prone to error when using a large database.
2. We present an R package (“optimos.prime”) that uses species’ abundance data and environmental data to calculate the optimum value and tolerance range of each species to each environmental factor, by weighted average. Additionally, the package can create caterpillar plots to show the results.
3. Using sample data from a phytoplankton database, we exemplify the use of the R package and its functions. A stand-alone version for Windows is also provided, and source code and documents are freely available on GitHub to encourage collaborative work.

## INTRODUCTION

When using organisms as indicators of environmental conditions, the key aspect to consider is the combination of environmental variables that are optimal for its existence, development, growth and reproduction (Verbitsky and Verbitskaya 2007). These aspects depend mostly on environmental factors, and the reproductive success of the species is related to an “ecological optimum” (Battarbee et al. 2010; ter Braak and Smilauer 1998). However, in natural ecosystems, the ecological optimum includes not only a single point value, but also the oscillations of the environmental variable around this value, and for each environmental variable there are lower and upper limits, below and above which the taxon is less likely to survive. These limits constitute the tolerance range of the taxon (Huggett 2004; Smith and Smith 2009; Cristóbal et al. 2014).

In this regard, the knowledge of autoecological data can be very helpful in the interpretation of the distribution, and presence/absence of different taxa in different environments (Clarke et al. 2006; Licursi et al. 2010). The most common approach used for its calculation is the weighted averaging, using the number of occurrences to adjust the tolerance assigned to each taxon to estimate optima and tolerance range in cases where taxa have unequal occurrences (Birks et al. 1990).

When employed, for instance, in water research and limnology, these methods have facilitated the reconstruction of past lake conditions (Oksanen et al. 1988; Birks et al. 1990; Bradshaw et al. 2002; Miettinen et al. 2005; Holden et al. 2008), the accurate estimation of contemporary limnological conditions in lentic and lotic environments (e.g: Potapova and Charles 2003; Meador et al. 2008; Yang et al. 2008; Licursi et al. 2010; Sathicq 2017), and the establishment of environmental quality standards for discerning the trends, causes, and consequences of watershed-land use (Cooper 1995; Kattel 2012; Arva et al. 2017). Quantitative autecological characteristics derived from small-scale regional datasets are useful for regional monitoring programmes, but since they are dependent on the restricted range and distribution of the environmental parameters in the dataset, they may not be appropriate for areas with different water chemistry characteristics. Reliable autecological data can be obtained only from datasets with large numbers of observations representing the full range of environmental conditions (Potapova and Charles 2003), and their calculation can be prone to errors, particularly when there is a significant number of species.

OPTIMOS PRIME is an open-source R script to facilitate the calculation of the ecological optima and tolerance ranges for each species to a given set of environmental variables. It is based on the weighted averaging procedure mentioned above. In this article we present its basic functions, providing an example of its use with data from phytoplankton samples that is available through the GitHub repository of the project. OPTIMOS PRIME is developed in R since it is increasing popularity as an analytical tool in the ecological sciences (e.g: Dixon 2003; Colchero et al. 2012; Metcalf et al. 2012; Revell 2012), although a stand-alone version for Windows is also available in the GitHub repository (https://github.com/limnolab/Optimos-Prime). Users can freely download a stable version of the package from the CRAN website (https://cran.r-project.org/web/packages/optimos.prime). Also, OPTIMOS PRIME development is being managed via GitHub to encourage collaborative development.

## DATA REQUIREMENTS AND INPUT

The calculation of species’ optima and tolerance range to environmental variables requires two input data frames (in the R terminology). The first data frame (“Environmental data”) contains all the values for all environmental parameters measured in the samples or sampling sites. The second data frame (“Species data”) contains the relative abundance of each species in each of the samples or sampling sites. Sample data frames are included in the package (/data subfolder).

### Environmental data

This data frame contains the environmental parameters measured (rows) by the sampled sites (columns). The first row has to contain the names of the sites, and the first column has to contain the names of the environmental parameters, or the code that the user will use as the default labels for the plots. A sample data frame is included in the package. Missing data should be left blank in the data frame.

### Species data

This data frame contains the relative abundance of each species (rows) in each sampled site (columns). The first row, as in the environmental data frame, has to contain the names of the sites, and the first column has to contain the names of the species, or the code that the user will use as the default labels for the plots.

These data frames can be already loaded in R or imported into R from a comma separated value file (CSV) created in a spreadsheet software (e.g. Microsoft Excel). If the data frames are not specified when the function is run, the user will be prompted to select a CSV file.

## CALCULATION OVERVIEW

There are three functions in the OPTIMOS PRIME package (Table 1). The function op_calculate() will use the environmental and species’ data frames to calculate the optima and tolerance range for each species to each environmental parameter. During the calculation, the environmental data needs to be transformed into log10; the op_calculate function assumes that the data is raw (not transformed), but if it is already transformed the parameter “islog10” (FALSE by default) of the function has to be set to TRUE.

**Table 1.**
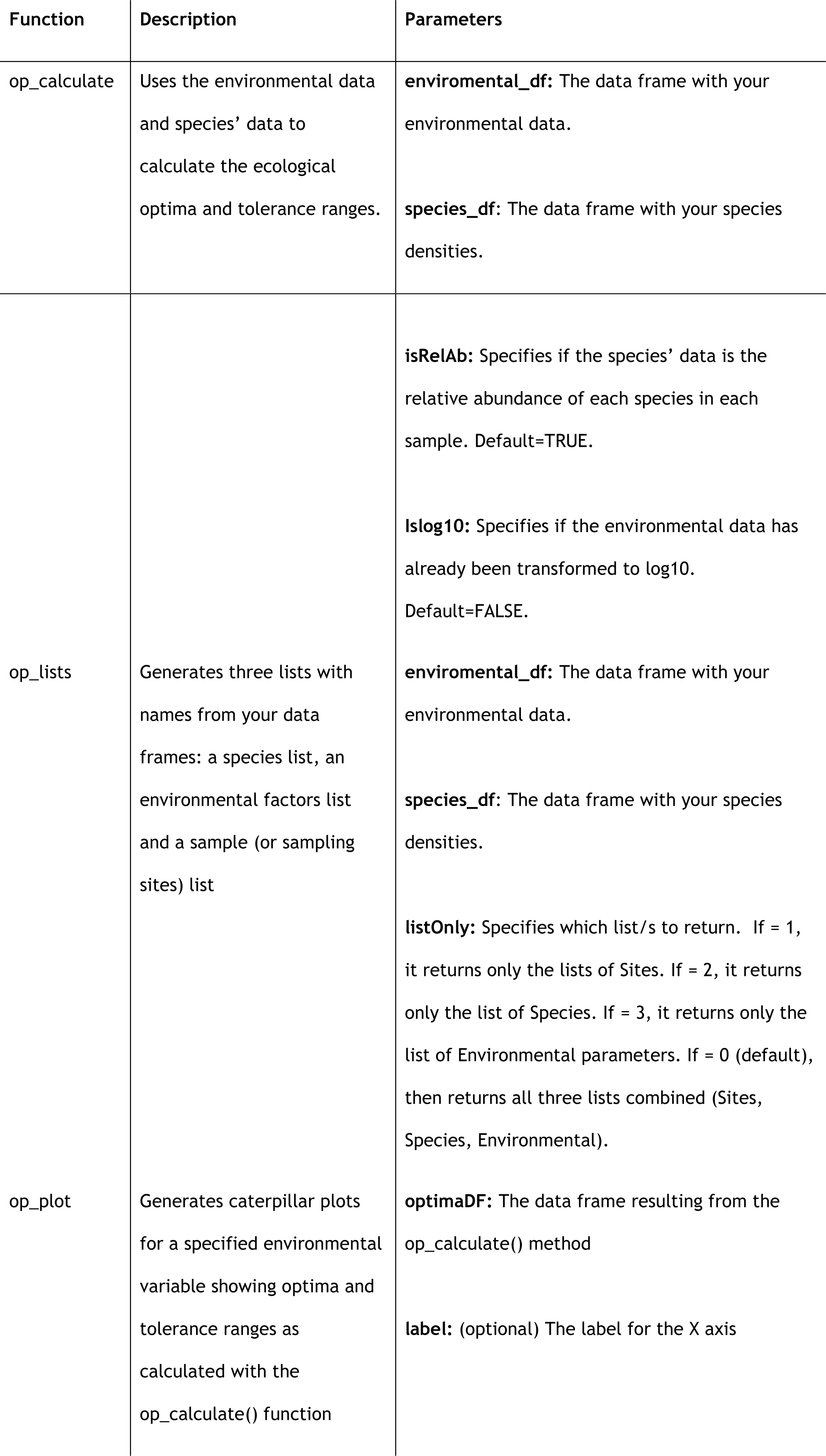
Functions included in the OPTIMOS PRIME package, their description and parameters.

The species’ data needs to be the relative abundance of each species in each sample. If this is not the case, and the data frame being used is the raw density of the species, then the “isRelAb” parameter needs to be changed to FALSE, and the function will convert it automatically to relative abundance values.

The weighted average estimates of the species optima (*u*_*k*_) is calculated as:

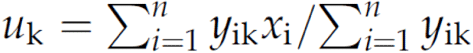

Where *y*_*ik*_ is the relative abundance of species k in sample *i*; *x*_*i*_ is the log10 value of the environmental parameter in sample *i*; and *n* is the total number of samples in the dataset. Tolerance, or weighted standard deviation (*t*_*k*_), is calculated as:

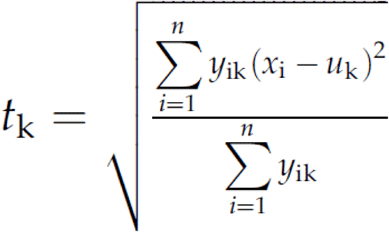

The op_lists() function allows the user to extract from the environmental and species data frames a list of the names of the sampling sites (Site list), of the names of the environmental variables (Environmental list), of the names of the species’ (Species list) or a single list including all the previous three lists. This can be used to check if the original data frames were constructed according to the specifications above, or to use the lists for further tests in R.

## PLOTTING RESULTS

The op_plot() function will provide with caterpillar plots for all species showing their optima and tolerance ranges to a certain chosen variable. This function uses the result data frame obtained from the op_calculate() function as input, provides the user with a choice to select which environmental variable to plot and draws an interactive caterpillar plot (Figure 1). This function depends on three other libraries that need to be installed along with the OPTIMOS PRIME package: *ggplot2, tidyverse*, and *plotly*. These three libraries should be automatically installed when Optimos Prime is loaded into R; otherwise they can be obtained from the CRAN repository.

**Fig 1.**
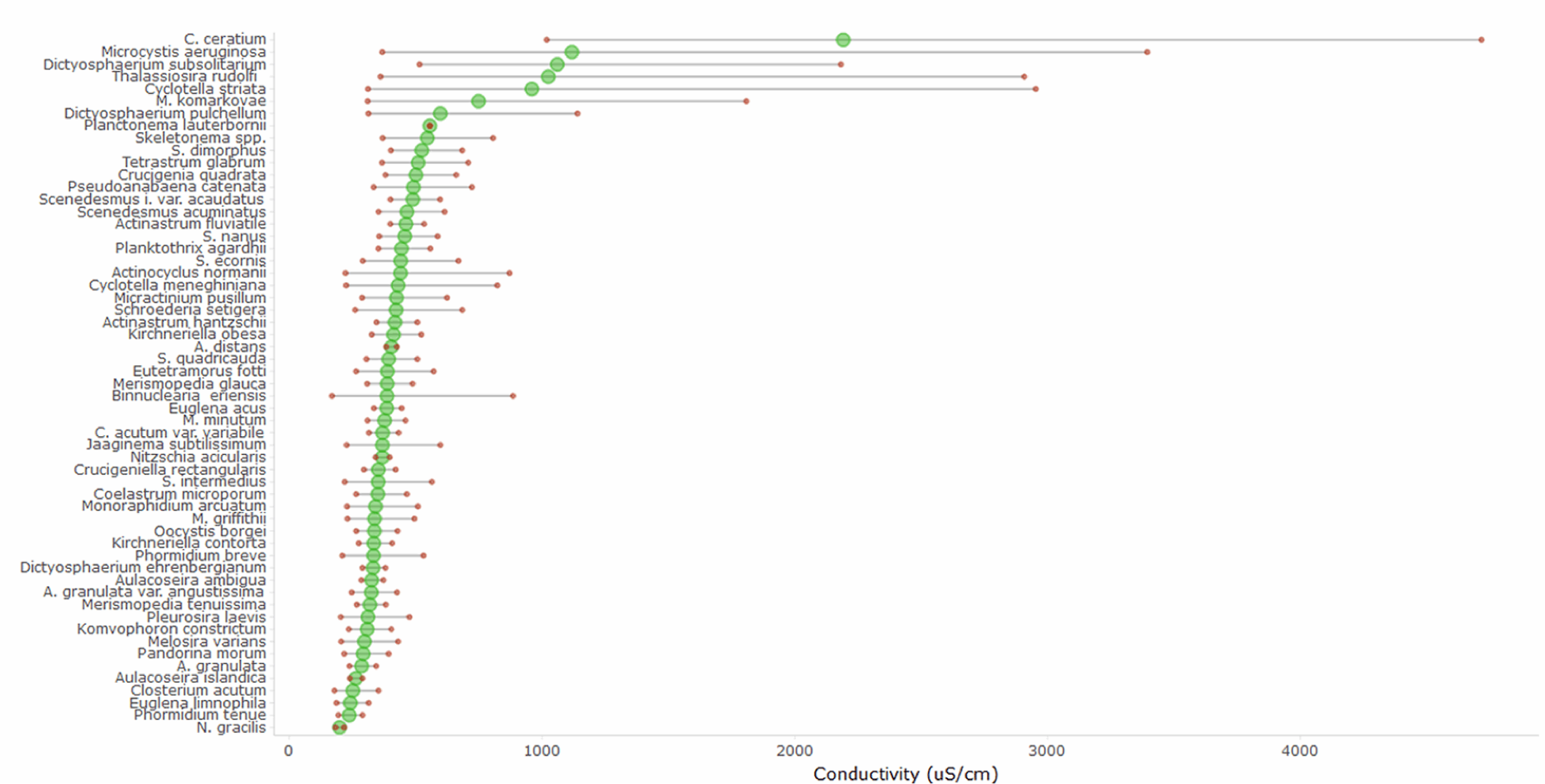
Caterpillar plot resulting from the op_plot() function with the example data, using “electrical conductivity” as an environmental factor.

## EXAMPLE APPLICATION

The following example uses two CSV files that are openly available in the GitHub repository of the project (https://github.com/limnolab/Optimos-Prime/tree/master/sample_data), although both data frames are also included in the package (/data subfolder). The file “environmental_data.csv” contains data from 11 environmental parameters (such as temperature, electrical conductivity, dissolved oxygen, etc.) in 50 samples. The file “species_data.csv” contains the relative abundance of 57 species of algal plankton in the same 50 samples. The data corresponds to a subset of the phytoplankton data used in Sathicq (2017).

For this example, we will assume that the user wants to load the data directly from the CSV file, and not from an already existing data frame in R. The following code will load the library and calculate the autoecological data using the input data frames, placing the results into a new data frame called “results_op”. All figures shown for this example are displayed if the user is using RStudio, although similar results will be obtained if using the R core.

~~~
> install.packages(“optimos.prime”)
> results_op <- op_calculate()
         “Select CSV matrices”
         [1] “Select ENVIRONMENTAL matrix”
         [2] “Select SPECIES matrix” “
         Calculating…”
         “Optimum values and tolerance range calculated and placed in the final data frame”
         “Use View() to view data frame with results”
~~~

Notice that no parameters were passed to the op_calculate() function, which resulted in the user being prompted to choose the CSV files from a dialog box in steps [1] and [2]. Also, the parameter islog10 was not set; its default value is FALSE, which assumes that the data is raw and was not previously transformed to log10.

The results of the analysis were placed in the data frame “results_op”, which can be viewed using View (results_op), as shown in Figure 2. Each row is a species, and each column consists of the ecological optima for each environmental parameter and its range of tolerance (columns HIGH and LOW for each parameter). If some species was absent from all sampling sites where the environmental variables were measured, the results show “NaN”, indicating missing data.

**Fig. 2.**
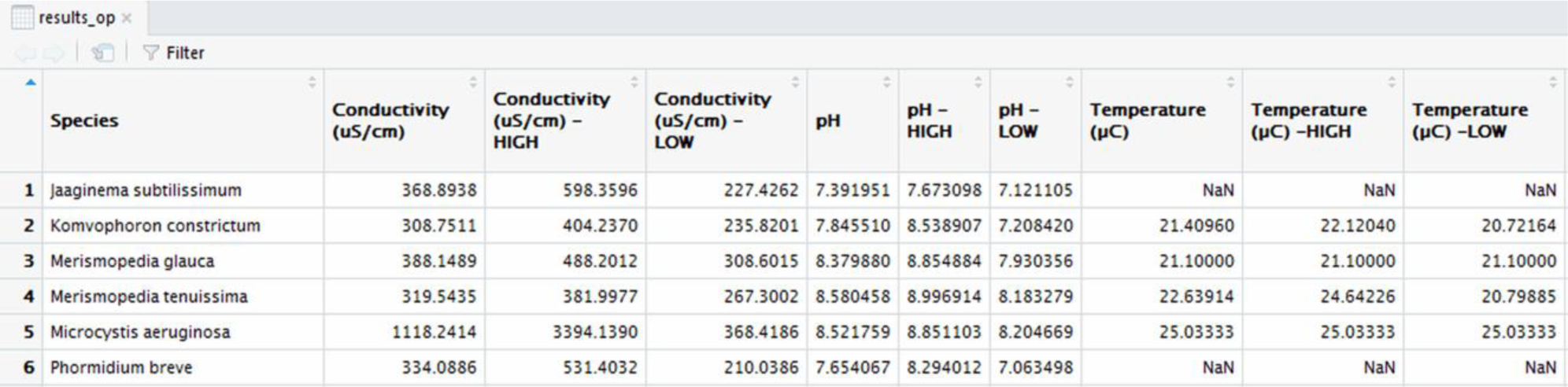
Sample of the resulting data frame from the optima and tolerance ranges calculation.

The caterpillar plots for (for example) the environmental parameter “Conductivity” can be now obtained by using the op_plot() function, specifying the recently obtained data frame.

~~~
> op_plot(results_op)
        1 Conductivity (uS/cm)
        2 pH
        3 Temperature (°C)
        4 Dissolved Oxygen (mg/L)
        5 BOD5 (mg/L)
        6 COD (mg/L)
        7 N-NO3 (mg/L)
        8 N-NO2 (mg/L)
        9 Turbidity(NTU)
        10 N-NH3 (mg/L)
        11 P-PO4 (mg/L)
> [Question] What variable number do you want to plot? 1
        “Plotting…”
        “Plot complete!”
~~~

After the function is run, a list of the environmental variables is shown in the terminal, and the user is prompted to type the number of the variable that is to be plotted (1 in this case is Conductivity). The interactive plot resulting is shown in Figure 1; green points mark each optimum and red points mark both high and low ends of the tolerance range for each species to that variable.

## WINDOWS STAND-ALONE VERSION

A stand-alone version for Windows is also available from the GitHub repository. The compressed file (.zip) includes an installer, and the software leads the user into loading the data frames in order. To begin the process, the user has to establish how many species, samples and environmental parameters is going to use; in the following tabs, the data frames can be pasted directly from the clipboard in its place (Figure 3). The output of the Windows version is similar to the results data frame from the R package described above, although it does not include a plotting function to create the caterpillar plots. The output can be exported back to CSV format, and the software is available both in English and Spanish.

**Fig 3.**
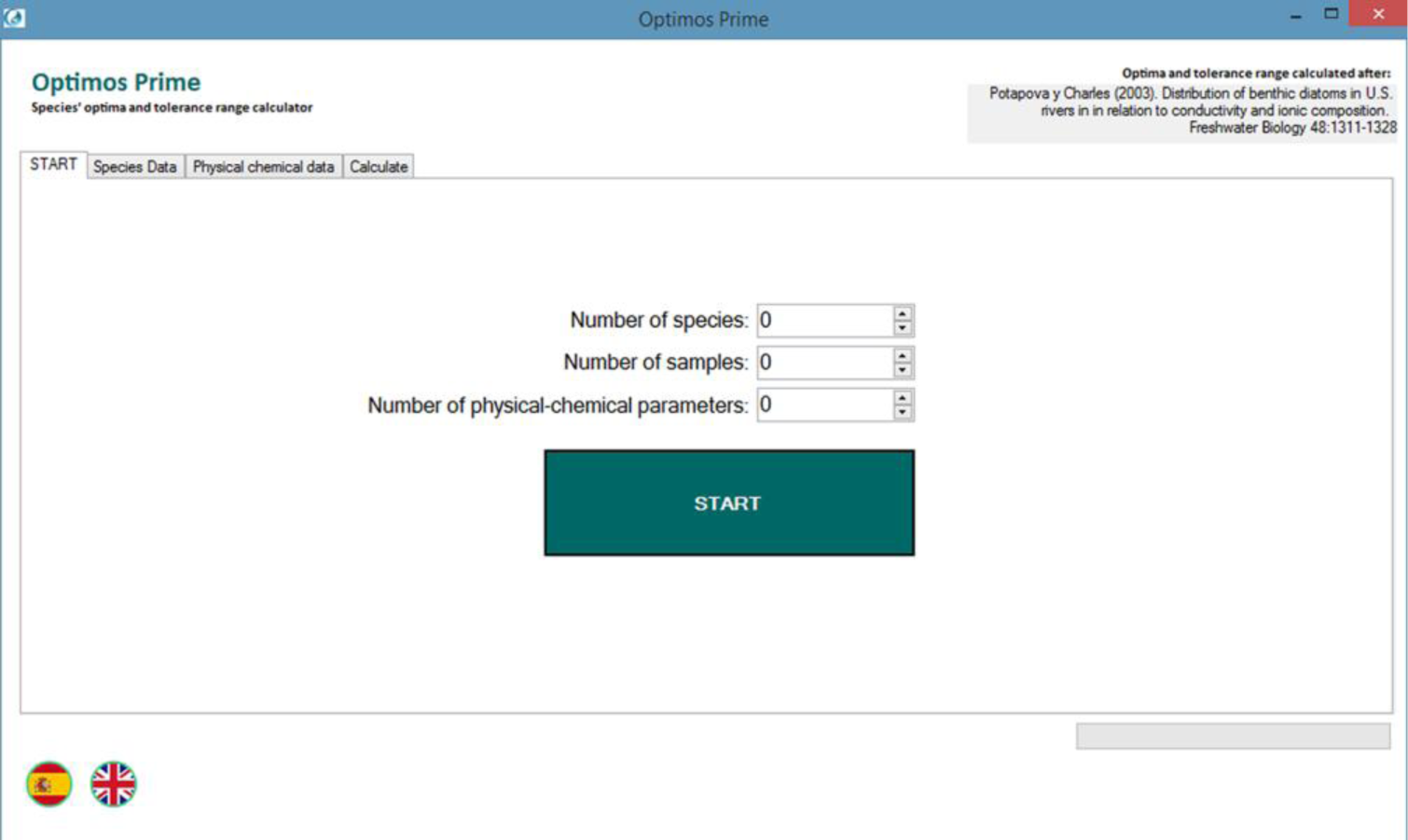
Main form of the OPTIMOS PRIME stand-alone version for Windows.

## CONCLUSIONS

We introduced the open-source R package OPTIMOS PRIME, which is designed to facilitate the calculation of the ecological optima and tolerance ranges for each species to a given set of environmental variables. The OPTIMOS PRIME package contains function for calculate optima and tolerance ranges, produce caterpillar plots, and generate lists with names from the data frames. In the GitHub repository of the package we provide the source code, along with two CSV files (“environmental_data.csv”, “species_data.csv”) with sample data, which users can download to test the package and build their data frames. A stand-alone Windows version is also available from the GitHub repository for non-R users. Future versions of OPTIMOS PRIME will include other algorithms for optima and tolerance range calculations, besides weighted averaging. Source code is publicly available through GitHub to encourage collaborative work.

## AUTHORS’ CONTRIBUTIONS

All authors contributed to software planning and manuscript writing. M.B.S. and M.M.N.G. provided sample data and software testing. J.C. led software development and coding.

## DATA ACCESSIBILITY

The OPTIMOS PRIME package can be downloaded from the Comprehensive R Archival Network (https://cran.r-project.org/web/packages/optimos.prime) or GitHub (https://github.com/limnolab/Optimos-Prime). A stand-alone Windows version can be downloaded from the repository as well. OPTIMOS PRIME is open source under GNU Public License.

## Notes

https://cran.r-project.org/web/packages/optimos.prime

